# Structure and function of virion RNA polymerase of crAss-like phage

**DOI:** 10.1101/2020.03.07.982082

**Authors:** Arina V. Drobysheva, Sofia A. Panafidina, Matvei V. Kolesnik, Evgeny I. Klimuk, Leonid Minakhin, Maria V. Yakunina, Sergei Borukhov, Emelie Nilsson, Karin Holmfeldt, Natalya Yutin, Kira S. Makarova, Eugene V. Koonin, Konstantin V. Severinov, Petr G. Leiman, Maria L. Sokolova

**Author notes:** Contributed equally (Arina V. Drobysheva, Sofia A. Panafidina).

## Abstract

CrAss-like phages are a recently described family-level group of viruses that includes the most abundant virus in the human gut^1,2^. Genomes of all crAss-like phages encode a large virion-packaged protein^2,3^ that contains a DFDxD sequence motif, which forms the catalytic site in cellular multisubunit RNA polymerases (RNAPs)^4^. Using *Cellulophaga baltica* crAss-like phage phi14:2 as a model system, we show that this protein is a novel DNA-dependent RNAP that is translocated into the host cell along with the phage DNA and transcribes early phage genes. We determined the crystal structure of this 2,180-residue enzyme in a self-inhibited, likely pre-virion-packaged state. This conformation is attained with the help of a Cleft-blocking domain that interacts with the active site motif and occupies the RNA-DNA hybrid binding grove. Structurally, phi14:2 RNAP is most similar to eukaryotic RNAPs involved in RNA interference^5,6^, although most of phi14:2 RNAP structure (nearly 1,600 residues) maps to a new region of protein folding space. Considering the structural similarity, we propose that eukaryal RNA interference polymerases take their origin in a phage, which parallels the emergence of the mitochondrial transcription apparatus^7^.

---

Transcription of bacterial, archaeal, and nuclear eukaryal genes is performed by multisubunit DNA-dependent RNA polymerases (RNAPs), complex molecular machines that have a common ancestor^4,8-10^. Their active site is located at the interface of two double-psi β-barrel (DPBB) domains that belong to two different polypeptide chains. One of the DPBB domains carries the universally conserved amino acid motif DFDGD, where the three aspartates coordinate Mg^2+^ ions required for catalysis^11,12^. Gene g066 of *Cellulophaga baltica* crAss-like phage phi14:2 encodes a 2,180-residue protein that shows a limited sequence similarity to one of the two DPBB domains of cellular RNAPs and contains a motif (^1361^DFDID^1365^) that is conserved in orthologs of this protein across the crAss-like phage family^2^. Gp66 protein has been identified as a component of the phage particle^3^. We hypothesized that gp66 is an evolutionarily divergent virion-packaged RNAP of phi14:2 that is delivered into the host cell early in the infection process where it transcribes the early phi14:2 genes. To test this hypothesis, we examined the *in vitro* and *in vivo* activity of gp66 and solved its crystal structure.

## RNAP gp66 transcribes single-stranded and denatured double-stranded DNA *in vitro*

We expressed recombinant gp66 in *Escherichia coli*, purified it (**Extended data Fig. 1**), and tested its RNA synthesis activity in a diverse set of assays.

First, we tested whether gp66 could extend the RNA primer of an 8-nucleotide long RNA-DNA hybrid in the presence of ribonucleoside triphosphates (rNTPs). This hybrid molecule mimics the nucleic acid structure in the transcription elongation complex^13^. Gp66 was inactive in this assay whereas both *E. coli* and T7 RNAPs extended the RNA primer (**Extended data Fig. 2**).

Next, we examined whether gp66 can initiate transcription of double-stranded and single-stranded DNA templates. Gp66 did not transcribe the genomic DNA of phage M13 in a double-stranded form and showed weak transcription of the phi14:2 genome (**Fig. 1a**). In contrast, single-stranded M13 genome and denatured phi14:2 DNA were transcribed very efficiently (**Fig. 1a, 1b**). The reaction products were resistant to DNase RQ1 treatment and sensitive to RNase T1 indicating that these high-molecular weight nucleic acids comprised entirely of newly synthesized RNA (**Fig. 1c**).

**Fig. 1.**
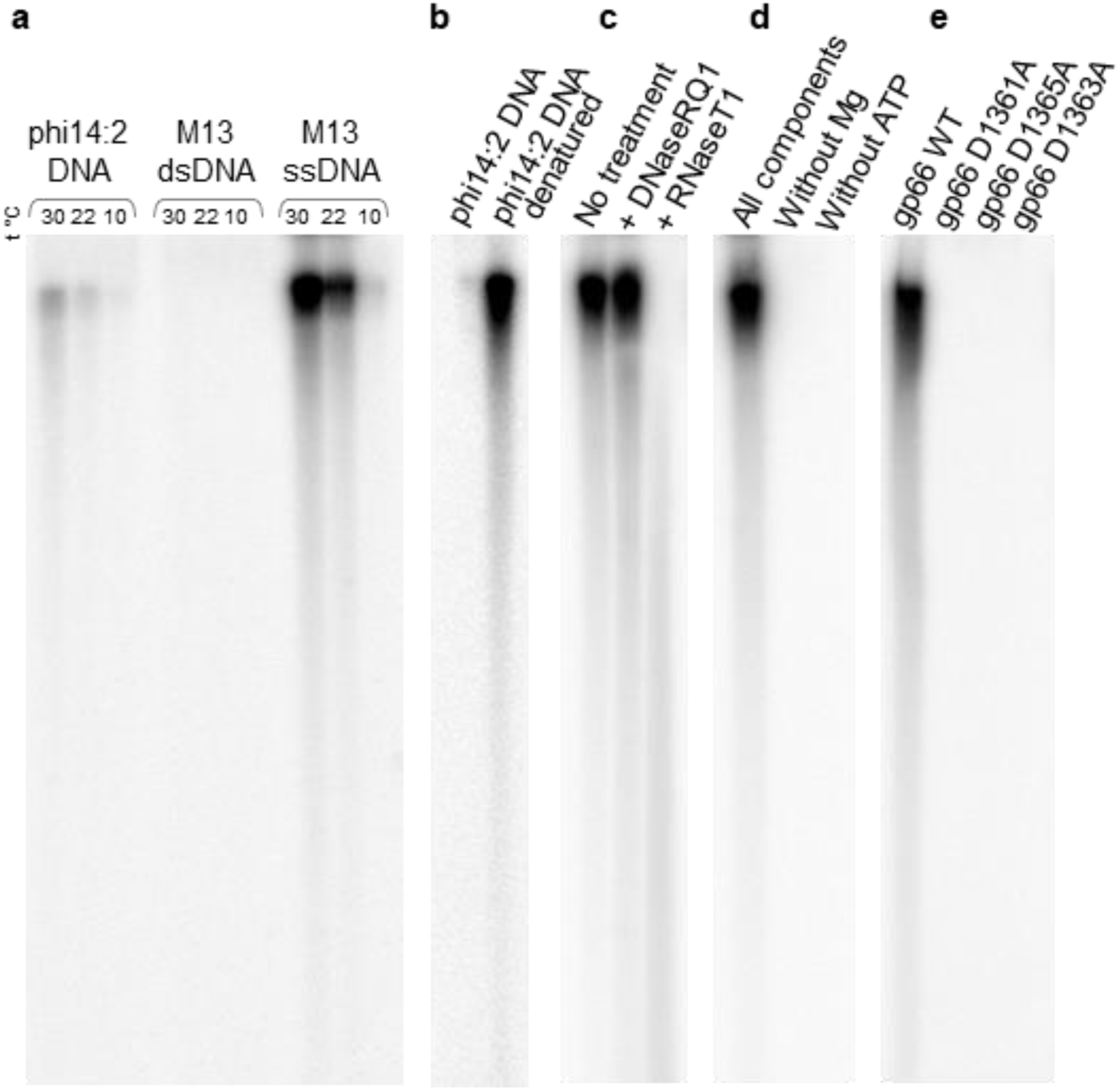
*In vitro* transcription activity of the phi14:2 RNAP gp66. **a**, Transcription by gp66 of genomic DNA of phages phi14:2 and M13 (double- and single-stranded forms) at 30, 22, and 10°C; the reaction products were resolved by electrophoresis in 5 % (w/v) denaturing 8 M urea polyacrylamide gel and revealed by autoradiography. **b**, Transcription by gp66 of native and denatured genomic DNA of phage phi14:2. **c**, Completed transcription reactions of phi14:2 genomic DNA were treated with DNase RQ1 or RNase T1 prior to loading on the gel. **d**, Activity of gp66 requires Mg ions and ATP. Denatured genomic DNA of phi14:2 phage has been used as a template. **e**, Transcription of phi14:2 denatured genomic DNA by wild-type gp66 and gp66 mutants carrying single alanine substitutions of each aspartate in the DFDID motif.

All experimentally characterized RNAPs require Mg^2+^ ions for template-dependent polymerization of rNTPs, and all three aspartates of the DFDGD motif must be present to form a Mg^2+^-binding site. Gp66 had no activity in the absence of Mg^2+^ or one of rNTPs (rATP, **Fig. 1d**). Furthermore, RNA synthesis activity of gp66 was abolished if any of the aspartates of the ^1361^DFDID^1365^ motif was replaced with an alanine (**Fig. 1e**).

## Transcription of phi14:2 genome during infection is organized in three temporal stages

Genes of phi14:2 can be divided into three classes according to the timing of transcript accumulation throughout the infection (**Fig. 2a,b, Supplementary Table 2**). These classes generally correspond to three functional modules – replicative, gene expression, and capsid genes – that have been identified by comparative genomics of crAss-like phages^2^. The early class includes the entire replicative gene module and is transcribed in the rightward direction (**Fig. 2a**). The middle and late classes are transcribed in the leftward direction and contain the gene expression and capsid modules, respectively (**Fig. 2a**). The gene expression module as well as other upstream middle genes are also actively transcribed late in infection because they encode putative virion proteins, namely, tail genes g071 and g072, the virion-packaged predicted RNAP gene (g066), and two neighboring genes (g065 and g067), whose products are also present in the phage phi14:2 particle^3^ (**Fig. 2a**).

**Fig. 2.**
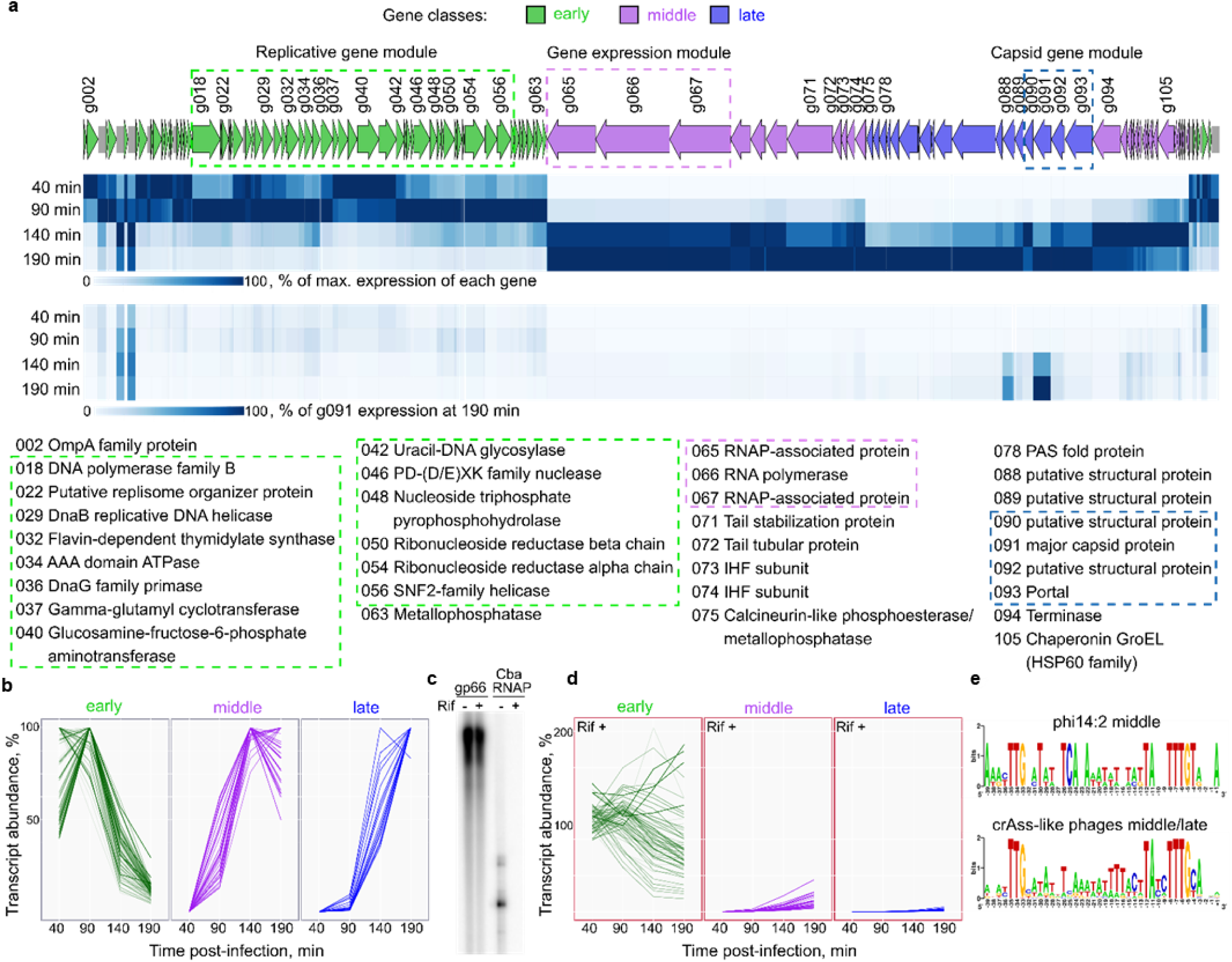
Global analysis of phi14:2 transcription during phi14:2 infection. **a**, Schematics of the phi14:2 genome^3^. ORFs are marked as arrows and numbered according to the study by Yutin et al^2^ (Supplementary Table 1). Intergenic regions larger than 50 base pairs are shown as grey rectangles. Replicative, gene expression, and capsid gene modules are marked by green, violet, and blue dashed frames, correspondingly^2^. Early, middle, and late genes are colored green, purple, and blue, correspondingly. Heat maps indicating the temporal pattern of phi14:2 transcripts and relative abundance of phi14:2 transcripts are shown below the genome. In the top heat map, the transcript abundance for each gene or intergenic region longer than 50 base pairs is normalized to the maximum transcript abundance for this particular gene/intergenic region; In the bottom heat map, the same quantities were normalized to the absolute maximum that corresponded to the abundance of the late gene g091 (major capsid protein) at 190 min post infection. **b**, Time courses of accumulation of individual phi14:2 transcripts divided into three temporal classes during infection; the y axis shows abundance of individual genes transcripts normalized to the maximal value for this gene obtained in Rif-libraries. **c**, Transcription by gp66 of denatured phi14:2 DNA and by *C. baltica* RNAP of a PCR-fragment containing the T7 A1 promoter in the absence and in the presence of rifampicin. **d**, Time courses of accumulation of early, middle, and late phi14:2 transcripts during infection in the presence of rifampicin; the y axis shows abundance of individual genes transcripts in Rif+ libraries normalized to maximal value for this gene obtained in Rif-libraries. **e**, WebLogos of phi14:2 middle promoters located upstream of middle genes g070, g075, g108, and g110 (top panel) and cumulative consensus of 126 crAss-like phage middle/late promoters (bottom panel, Supplementary file 1).

The transcript of the major capsid protein gene (g091) was the most abundant (**Fig. 2a**). Remarkably, two long intergenic regions (g004-g005 and g005-g006 junctions) were transcribed at a higher level than most protein coding genes (**Fig. 2a, Supplementary Table 2,3**). These intergenic regions are transcribed in the rightward direction and are present at the earliest time point sampled (40 min post-infection), but in contrast to all other early genes transcripts, their abundance does not drop later in infection (**Fig. 2a**). The functions of these long non-coding RNAs remain to be determined.

## Virion-packaged gp66 transcribes early genes of phi14:2

*In vitro*, gp66-dependent transcription was resistant to rifampicin, an inhibitor of bacterial RNAPs^14^, whereas *C. baltica* RNAP-dependent transcription was sensitive (**Fig. 2c**). This finding made it possible to examine the role of gp66 and *C. baltica* RNAP in the transcription of phi14:2 genome *in vivo*. Addition of rifampicin to a *C. baltica* culture infected with phi14:2 increased the relative abundance of phi14:2 transcripts reads in libraries obtained for every time point sampled and, conversely, reduced the abundance of *C. baltica* reads (**Extended data Fig. 3**). Rifampicin severely inhibited the transcription of the middle and late genes of phi14:2, whereas the transcription of the early genes was only moderately affected (**Fig. 2d** in comparison with **Fig. 2b**). Thus, the early genes of phi14:2 are transcribed by gp66, a rifampicin-resistant RNAP, which must be translocated into the host alongside phi14:2 DNA.

## Middle and late genes of phi14:2 are transcribed by the host RNAP

In order to delineate the 5’ends of phi14:2 transcripts and identify promoters, we performed primer extension analysis of RNA purified from infected cells (**Extended data Fig. 4**). Early transcripts that are synthetized by gp66 RNAP did not contain detectable common upstream motifs. By contrast, an extended tripartite motif was present upstream of middle genes transcripts (**Extended data Fig. 4, Fig. 2e)**. Two blocks of this motif resembled the ‘−35’and ‘−10’promoter consensus elements recognized by bacterial RNAPs containing primary σ-factors^15^. Indeed, *E. coli* σ^70^-RNAP holoenzyme transcribed PCR fragments with such promoters *in vitro* (**Extended data Fig. 4d**). The 5’ends of these transcripts matched those of RNAs purified from infected *C. baltica* cells (**Extended data Fig. 4**).

The genomes of crAss-like phages in the candidate genus VI of the beta-crassvirinae subfamily^16^ possess motifs that are similar to phi14:2 middle promoters (**Supplementary file 1, Fig. 2e**). These putative promoters are located upstream of homologs of phi14:2 middle and late genes (**Supplementary file 1**). Thus, middle and late genes in other crAss-like phages are likely also transcribed by the respective host RNAPs.

## Crystal structure of gp66 reveals a unique active site conformation

To better characterize gp66 RNAP, we crystallized it and solved its structure to a resolution of 3.5 Å. Two different crystal forms were produced (monoclinic and orthorhombic) and both contained two RNAP molecules in the asymmetric unit (4,388 amino acids including affinity tags). The phase information was obtained with the help of Ta_6_Br_12_ and SeMet derivatives by the single wavelength anomalous diffraction technique. The atomic model comprising 2,166 amino acids was refined to R/R_free_ values of 0.19/0.24 and contained 0.02% Ramachandran outliers (**Table 1**).

**Table 1.** Data collection and refinement statistics.

|  | Ta<sub>6</sub>Br<sub>12</sub> soaked | SeMet |
| --- | --- | --- |
| **Data collection** |  |  |
| Space group | P2<sub>1</sub>2<sub>1</sub>2 | P2<sub>1</sub>2<sub>1</sub>2 |
| Cell dimensions |  |  |
| *a*, *b*, *c* (Å) | 270.121, 299.343, 93.402 | 266.441, 297.181, 92.015 |
| α, β, γ (°) | 90.00, 90.00, 90.00 | 90.00, 90.00, 90.00 |
| Resolution (Å) | 50.0 – 3.75 (3.99 – 3.75) * | 50.0 – 3.50 (3.71 – 3.50) |
| *R*<sub>merge</sub> | 0.151 (1.450) | 0.176 (0.942) |
| *I* / σ*I* | 13.04 (1.60) | 11.90 (2.25) |
| Completeness (%) | 98.7 (94.3) | 99.7 (99.4) |
| Redundancy | 13.85 (13.20) | 7.90 (7.90) |
| **Refinement** |  |  |
| Software (version) |  | Phenix.refine (1.41) |
| Resolution (Å) |  | 50.0 – 3.50 |
| No. reflections |  | 177,613 |
| *R*<sub>work</sub> / *R*<sub>free</sub> |  | 0.191 / 0.244 |
| No. atoms |  |  |
| Protein |  | 34,688 |
| Ligand/ion |  | 26 |
| Water |  | 0 |
| *B*-factors |  |  |
| Protein |  | 88.81 |
| Ligand/ion |  | 72.90 |
| R.m.s. deviations |  |  |
| Bond lengths (Å) |  | 0.003 |
| Bond angles (°) |  | 0.568 |

\*Values in parentheses are for highest-resolution shell.

The structure of gp66 is most similar to that of *Neurospora crassa* single-subunit DNA/RNA-dependent RNAP QDE-1. QDE-1 and its homologs in other eukaryotes synthesize short RNAs involved in RNA interference^5,6^. Both gp66 and QDE-1 contain two DPBB domains that belong to two different subunits in multisubunit RNAPs (β and β’ in bacterial RNAP) within a single chain. Furthermore, in both gp66 and QDE-1 the two DPBB domains are connected by a similar ∼140-residue long Connector domain (residues 1095 – 1238 and 793 – 919 in gp66 and QDE-1, respectively) (**Fig. 3**).

**Fig. 3.**
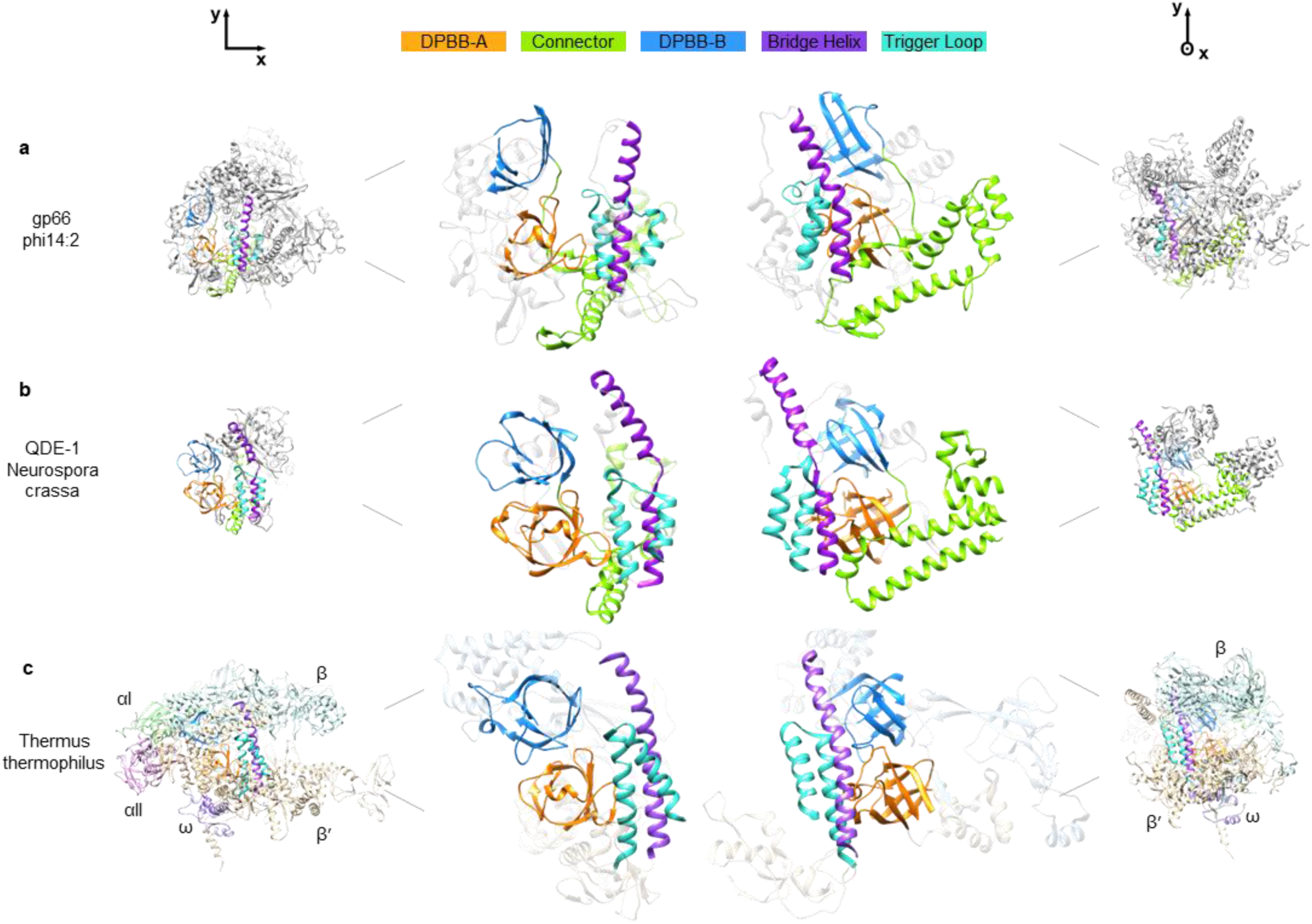
Phi14:2 RNAP gp66 is related to single- and multi-subunit RNAPs. **a, b**, and **c**, Crystal structures of phi14:2 gp66, QDE-1 from *N. crassa* (PDB 2J7N^6^), and *T. thermophilus* RNAP (PDB ID 2O5J^17^) are shown as ribbon diagrams, respectively. Conserved structural elements are colored according to the color code given above the top panels. Dissimilar domains of QDE-1 and gp66 are shown in gray color. Each of the five subunits comprising the T. thermophilus RNAP (αI, αII, β, β′, and ω) is rendered in a distinct color.

Gp66 RNAP shares two conserved structural elements and, possibly, corresponding functional features with multisubunit cellular RNAPs. Specifically, gp66 contains a trigger loop (residues 1598 – 1636), which loads rNTPs into the active site in multisubunit RNAPs^17,18^, and a bridge helix (residues 1529 – 1559) that is essential for RNAP translocation along the template^19-21^ (**Fig. 3**). Both, the trigger loop and the bridge helix of gp66 are more similar to those of QDE-1 than to the corresponding elements of cellular RNAPs. All strictly conserved residues of gp66 are located around the ^1361^DFDID^1365^ motif and most of them have counterparts in QDE-1 and/or multisubunit RNAPs (**Extended data Table 1**).

The overall structural similarity of gp66 to QDE-1 and multisubunit RNAPs is, however, low. Automatic superposition^22^ of gp66 onto QDE-1 identifies 479 equivalent residues that display 8.6% sequence identity and whose Cα atoms superimpose with a root mean square deviation (RMSD) of 3.5 Å. Superposition of gp66 onto *T. thermophilus* RNAP contains 489 residues with 8.2% sequence identity and an RMSD of 4.2 Å.

The structure of gp66 presents multiple unique features. Besides the two DPBB domains, Connector, and two structural elements involved in catalysis (the trigger loop and the bridge helix), the rest of gp66 domains comprising nearly 1,600 residues have no homologs that could be identified with existing tools^23^. The functions of these domains remain to be determined. Furthermore, the ^1361^DFDID^1365^ catalytic motif of gp66 is in a conformation that is incompatible with catalysis (**Fig. 4**). In all previously studied RNAPs, the fourth position in this motif is occupied by a glycine, and the three aspartate side chains point roughly towards the same point where they coordinate a Mg^2+^ ion required for catalysis. As shown above, Mg^2+^ and each of the aspartates of the catalytic motif are required for gp66 RNA synthesis activity, so this motif must be responsible for catalysis despite its unusual conformation in the crystal structure. Thus, in an actively transcribing gp66 RNAP, the catalytic motif apparently refolds to allow Mg^2+^ ion coordination by the three aspartate side chains. One way to accomplish this is for the isoleucine to adopt a left-handed turn conformation. RNAPs in all crAss-like phages contain an isoleucine or valine in the fourth position of the catalytic motif, so their active sites conformations and properties are likely to be similar to those of gp66.

**Fig. 4.**
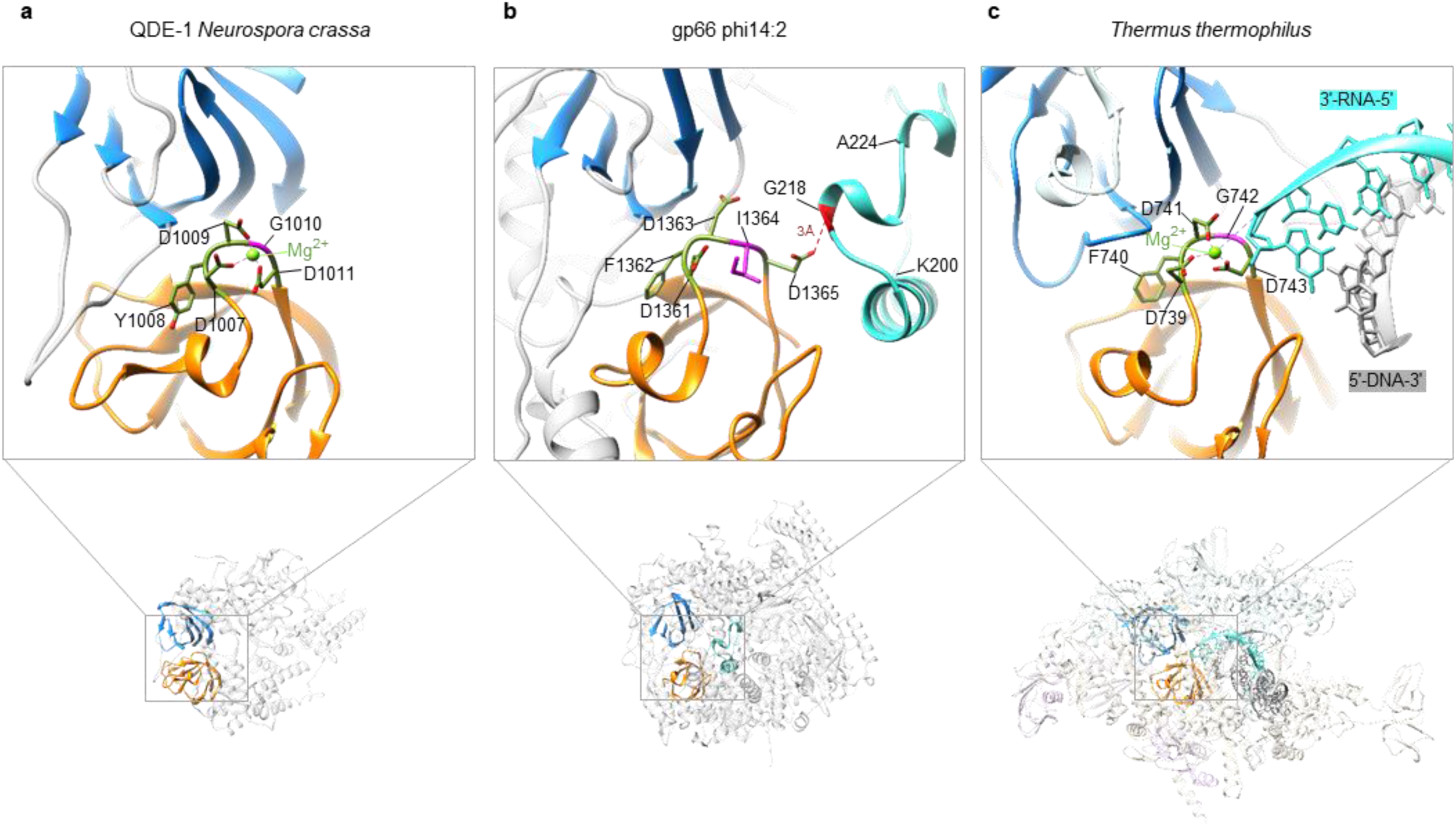
Cleft-blocking domain occupies the RNA-DNA hybrid binding site in phi14:2 RNAP gp66. **a, b**, and **c**, The structure of the active site of QDE-1 from *N. crassa* (PDB 2J7N^6^), phi14:2 gp66, and *T. thermophilus* RNAP (PDB ID 2O5J^17^), respectively. The active site of phi14:2 RNAP gp66 (**b**) is in a conformation incompatible with Mg binding.

## Regulation of activity of virion RNAPs of crAss-like phages

A notable feature of the gp66 structure is that its RNA-DNA hybrid binding cavity is occupied by a Cleft-blocking domain (residues 196 – 233) (**Fig. 4**). Besides forming a number of interactions with the cavity ‘walls’, it interacts with the catalytic ^1361^DFDID^1365^ motif (there is a nearly ideal hydrogen bond between G218 and the side chain of D1365) (**Fig. 4**). This interaction stabilizes the unusual conformation of the catalytic motif.

We hypothesize that the crystal structure represents a self-inhibited, pre-virion-packaged form of the enzyme as is likely required by the virion assembly pathway. At late stages of infection, newly synthesized copies of gp66 have to be available for packaging into the virus particle and thus have to be excluded from transcription of the phage genome. Gp66 attains its fully active conformation upon translocation into the cell during infection. In the active conformation, the Cleft-blocking domain and, possibly, the domain upstream of it, refold or are cleaved to free RNA-DNA hybrid binding grove. Notably, recombinant gp66 shows a strong *in vitro* single-stranded DNA transcription activity (**Fig. 1a,b**). Most likely, a single-stranded DNA template fits into the remaining space in the cleft and is able to displace the Cleft-blocking domain from the cavity for transcription to take place.

The self-inhibited conformation of virion-packaged gp66 RNAP parallels the assembly-coupled maturation in other viruses. This process is typically accompanied by a large-scale conformational change of the virus particle and involves proteolysis^24-26^. Whether activation of gp66 RNAP requires proteolysis or is accomplished by a novel and unique mechanism, remains to be determined. Notably, orthologs of gp66 RNAP and two proteins encoded by the adjacent genes (g065 and g067 in phi14:2) are present in the virions of phi14:2^3^ and other crAss-like phages^27^. In some phages, the three proteins are fused into a single huge polyprotein^2^, which is likely cleaved into individual components (one of which is the RNAP). One of these proteins (gp65 of phi14:2 and its orthologs) contains a Zincin-like metal-dependent protease domain^2^ that might be involved in the activation of RNAP and/or functions to digest the host peptidoglycan layer.

## Eukaryotic RNAPs involved in RNA interference and crAss-like phage RNAP share a common ancestor

QDE-1 and its orthologs comprise a family of RNAPs that is widespread in eukaryotes and is likely to have been present in the Last Eukaryotic Common Ancestor (LECA)^28,29^. These proteins were originally characterized as RNA-dependent RNAPs and were directly implicated in the production and/or amplification of small interfering RNAs^5^. However, it has been subsequently shown that *in vitro* they transcribe single-stranded DNA much more robustly than RNA^30,31^. Moreover, Replication Protein A (a single-stranded DNA-binding complex) and DNA helicase QDE-3 are required for RNA synthesis activity of QDE-1 on single-stranded DNA templates and for RNA silencing^31^. Thus, synthesis of small interfering RNAs by QDE-1 and related enzymes likely begins from transcription of a DNA template.

Structural similarity of crAss-like phage and QDE-1 RNAPs and the critical role of DNA-binding proteins in the function of the latter^31^, strongly suggests that the RNA interference RNAP was acquired by the LECA (or an earlier organism) from a phage, which infected a protomitochondrial endosymbiont^29^. This evolutionary scenario mimics the accepted view of the emergence of the mitochondrial transcription apparatus that takes its origin in an unrelated single-subunit RNAP of a T7-like phage^7^.

## Materials and Methods

### Bacterial and phage growth conditions, biological properties of phi14:2

*Cellulophaga* phage phi14:2 and its host *Cellulophaga baltica* strain #14 were previously isolated^32^. *C. baltica* strain #14^32^ was grown at room temperature (RT) on agar plates (12 g sea salt (Sigma), 1 g yeast extract (Helicon), 5 g Bacto Peptone (Helicon), and 15 g of agar (Helicon) per liter). Bacterial colonies were visible after 2-3 days of incubation. A single colony was inoculated into MLB liquid media (12 g sea salt (Sigma), 0.5 g yeast extract (Helicon), 0.5 g Bacto Peptone (Helicon), 0.5 g casamino acids (Difco), 3 mL glycerol (Sigma) per liter) and grown without agitation overnight. A high titer phi14:2 lysate was prepared using the top-agar plating technique as follows: 100 μL of the phi14:2 phage lysate diluted in MSM buffer (450 mM NaCl (Helicon), 50 mM MgSO_4_ (Panreac), 50 mM Tris-HCl (Sigma), pH 8.0, 0.01% gelatin (Dr.Oetker)) was mixed with 300 μL of bacterial overnight culture and 5 mL of molten soft agar (MSM buffer containing 0.4% TopVision Low Melting Point Agarose (ThermoFisher Scientific)) cooled to 32°C; the suspension was dispersed on agar plates. Plates were incubated at RT in the dark overnight. Further, 4 mL of MSM buffer was added to fully lysed plates, the top-agar surface was shredded and the plates were shaken for 30 min at RT, the liquid suspension was collected and centrifuged (4°C, 10,000 g, 10 min). The supernatant was 0.22-μm filtered (PES membrane filters, BIOFIL). The resultant phage stock (∼ 10^10^ - 10^11^ PFU/mL) was stored at 4°C.

To plot the growth curves of the *C. baltica* during infection by phi14:2, *C. baltica* cultures (n=3) were infected at OD_600_∼0.11 with phi14:2 at different multiplicity of infection (MOI) levels (0.01, 0.1, 1 and 10). The growth was monitored using EnSpire Multimode Plate Reader (PerkinElmer) by measuring OD every 30 min during 48 hours. At MOI of 10, culture lysis was observed 3.5 hours post-infection (**Extended data Fig. 5a**)

To perform single-burst experiment, the *C. baltica* cultures (n=3) were infected at OD_600_∼0.15 with phi14:2 at a MOI of 0.5 and immediately split into two flasks; one of the cultures was supplemented with rifampicin (10 μg/mL). Aliquots of infected cultures were withdrawn every hour. The number of plaque forming units (PFU) was determined by the top-agar plating technique. The latent period was 3 hours (**Extended data Fig. 5b**). During the next 3 hours, a gradual, ∼ 20-fold, increase of the number of plaque forming units in the culture was observed (**Extended data Fig. 5b**). Addition of rifampicin – an inhibitor of bacterial RNA polymerase (RNAP) – prevented the production of phage progeny (**Extended data Fig. 5b**).

### *C. baltica* genome sequencing and assembly

Genomic DNA of *C. baltica* strain #14^32^ was extracted from 2 mL of overnight culture by Genomic DNA Purification Kit (Thermo Fisher Scientific) according to manufacturer’s protocol for Gram-negative bacteria. DNA libraries were generated by the Skoltech Genomics Core Facility using NEBNext Ultra II DNA Library Prep Kit for Illumina (NEB) following the manufacturer’s instructions and sequenced on Miseq (Illumina) instrument using Miseq reagents v.3, 600 cycles. Sequence reads were quality-checked using FastQC v0.11.8. The adapters and low-quality sequences were eliminated using Trimmomatic v0.38. Reads were assembled by SPAdes v3.13.0 with standard parameters.

### Sample collection and RNA purification for RNA-Sequencing

*C. baltica* strain #14 culture was grown to OD_600_ of 0.14, split into two flasks and one of the two cultures was supplemented with rifampicin (10 μg/mL). The cultures were infected with phi14:2 at a MOI of 10. To synchronize the infection, 40 min after the infection, the two cultures were centrifuged (RT, 5000g, 15min) and the pellets were resuspended in the same amount of fresh MLB medium with and without rifampicin correspondingly. At various time points (40, 90, 140, 190 min post-infection), 20-mL aliquots of infected cultures were withdrawn, collected by centrifugation and kept at −20°C. Efficiency of infection was measured by comparing colony-forming units (CFU) before the infection with CFU determined 90 min post-infection. Total RNA was purified from the cell pellets using GeneJET RNA Purification Kit (Thermo Fisher Scientific) following the manufacturer’s instruction (Bacteria Total RNA Purification Protocol) with an additional step: after resuspension in the Lysis Buffer the cells were disrupted by sonication (two rounds of exposure for 10 seconds with a 50 seconds interval at an amplitude 20% (Q500 Sonicator by Qsonica)). RNA samples (5 μg of each) were treated with RNase-free DNase I (Thermo Fisher Scientific) in the presence of RiboLock (Thermo Fisher Scientific) for 1 h at 37°C and RNA was subsequently purified by GeneJET RNA Purification Kit (Thermo Fisher Scientific) according to the manufacturer’s instructions. RNA concentrations were determined with a NanoDrop spectrophotometer. The overall levels of rRNA did not change throughout the infection, as determined by visual inspection of agarose gel lanes.

### RNA-Seq library preparations and sequencing

cDNA libraries were constructed by the Skoltech Genomics Core Facility as follows. Ribosomal RNA was depleted from the total-RNA samples using Ribo-Zero rRNA Removal Kit (Illumina), according to the manufacturer’s protocol. Subsequently, strand-specific cDNA libraries were generated by NEBNext Ultra II Directional RNA Library Prep Kit for Illumina (NEB) following the manufacturer’s instructions, with exception of fragmentation time (10 minutes instead of 15). Eight libraries were created (40 min Rif-, 90 min Rif-, 140 min Rif-, 190 min Rif-, 40 min Rif+, 90 min Rif+, 140 min Rif+, 190 min Rif+). The single-end strand-specific sequencing with 84 bp length of the reads was performed on an Illumina Nextseq500. In total, 13 million to 25 million sequence reads were obtained for each cDNA library.

### RNA-Seq data analysis

The raw reads were subjected to quality filtering and adaptor trimming using Trimmomatic v0.38^33^ with the following parameters: SE-phred33 Illuminaclip:TruSeq3-se:2:30:10 leading:3 trailing:3 slidingwindow:4:15 minlen:36. The quality before and after processing was examined using FastQC tool. Processed reads were mapped to the reference sequences (phi14:2 genome (NC_021806.1) and the *C. baltica* strain #14 genome (BioProject ID PRJNA552277) using bowtie2 v2.3.4.3 with default settings. Overall, 88 – 99 % of reads from each library aligned with the reference genomes of *C. baltica* and phi14:2 in a strand-specific manner. Ratio of phage and host transcripts abundances is shown in **Extended data Fig. 3**. The quantification of reads by phage genes was performed using featureCounts function from the Rsubread package v1.34.3 in a strand-specific mode and allowed multiple overlapping of reads with features; other parameters were set to default. RPKM (Reads Per Kilobase of transcript, per Million mapped reads) values were calculated with normalization on a total number of mapped reads (**Supplementary Tables 2,3**). These RPKM values were used to create the abundance curves and heat maps (**Fig. 2**).

### Criteria for classification of phi14:2 genes

Each gene was assigned to one of three temporal classes – Early, Middle, or Late – according to its transcript abundance within a certain period post infection. The dynamics of transcript abundance was quantified with the help of a Log-Fold Change parameter (LogFC) that was calculated as follows: LogFC_XvsY = log_10_A(Y) - log_10_A(X), where A(X) and A(Y) are normalized transcript abundances of the gene at time points X and Y post infection (**Supplementary Table 2**).

The maximum value of transcript abundance of the Early class genes was within the first 90 min post infection, so their LogFC values obeyed the following criterion: LogFC_90vs140 < 0 and LogFC_140vs190 < 0. The maximum value of transcript abundance of the Middle class genes was in the 90-190 min post infection period and the increase of abundance within that period did not exceed 10 times: LogFC_90vs190 > 0 and LogFC_90vs190 ≤ 1. The transcript abundances of the Late class genes increased by more than 10 times in the 90-190 min post infection period: LogFC_90vs190 > 1.

### RT-qPCR

Results obtained by RNA-Seq for both Rif- and Rif+ cultures were validated by reverse transcription-quantitative PCR (RT-qPCR) with primers specific to randomly chosen Early, Middle and Late phage genes (**Supplementary Table 5** and **Extended data Fig. 6**).

Total RNA was purified as described in the sample collection and RNA purification section. First-strand cDNA synthesis was performed with Maxima reverse transcriptase (Thermo Fisher Scientific) and random hexamer primers (Thermo Fisher Scientific) with 150 ng of total RNA according to the manufacturer’s instructions. The subsequent qPCR analysis was performed using iTaq Universal SYBR Green Supermix (Bio-Rad), on Applied Biosystems QuantStudio 3 amplifier with primers listed in **Supplementary Table 5**. The cycle threshold (Ct) values of the 16S RNA were used to normalize the Ct values of selected phi14:2 transcripts (ΔCt = (mean Ct gene) – (mean Ct 16S rRNA)). To follow the relative differences in amplicon concentrations for different samples, a 2^(−ΔCt) value was used.

### Primer extension and sequencing reactions

Gene-specific primers (**Supplementary Table 4**) were labeled with [γ-^32^P]ATP by phage T4 polynucleotide kinase (New England Biolabs), as recommended by the manufacturer. Primer extension reactions were performed with 1 pmol of [γ-^32^P]ATP end-labeled primers and 5 μg of total RNA using Maxima reverse transcriptase (Thermo Fisher Scientific) according to the manufacturer’s instructions. Reactions were terminated by the addition of an equal volume of denaturing loading buffer (95% formamide, 18 mM EDTA, 0.25% SDS, 0.025% xylene cyanol, 0.025% bromphenol blue). Sequencing reactions were performed with the same primers as the ones used for the primer extension reactions and with PCR fragments (amplified from phi14:2 genomic DNA) using the USB Thermo Sequenase Cycle Sequencing Kit (Thermo Fisher Scientific) according to manufacturer’s instructions. The reactions were terminated as above. The reaction products were resolved on 6–8% (w/v) denaturing polyacrylamide gels and visualized with Typhoon FLA scanner (GE Healthcare). In total, 23 phi14:2 genome regions, which could contain promoters were analyzed and primer extension products for ten of them were detected (**Supplementary Table 4**).

### Search for nucleotide sequence motifs

To identify motifs similar to the phi14:2 Middle promoter motif in genomes of other crAss-like phages, 36 previously analyzed representative genomes^2^ (ftp://ftp.ncbi.nih.gov/pub/yutinn/crassphage_2017/), the 242 genomes from a subsequent study^16^ and the genome of phicrAss001^27^ were scanned. First, we searched for occurrences of the phi14:2 Middle promoter motif by using the program FIMO^34^ (Supplementary file 1). Thirteen genomes that contained at least four unique hits with a score greater than 1 and genome of IAS phage that contained three unique hits were used to create new consensus motifs, which were then used as new templates to search motifs in the same 14 genomes. The new searches resulted in 137 hits of which 17 corresponded to coding regions and the rest to intergenic regions. All but four intergenic hits were in the sense direction.

Identification of coding regions required annotation of the following twelve phage genomes: cs_ms_27, err843924_ms_3, ERR844029_ms, ERR844058_ms_2, ERR844065_ms_1, SRR4295173_s_14, SRR4295175_s_4, eld298-t0_s_3, ERR844030_ms_2, cs_ms_22, Fferm_ms_11, and HvCF_E4_ms_5. HMM profiles of conserved protein families of crAss-like phages from Yutin et al^2^ were generated from multiple sequence alignments published by Yutin et al^2^ using hmmbuild tool from the HMMER v3.1b2 package (http://hmmer.org/) with default settings. tRNA and tmRNA genes were predicted using ARAGORN v1.2.38^35^. ORFs were predicted with Prodigal v2.6.3^36^. Amino acid sequences of predicted ORFs were scanned against Pfam-A v32.0 supplemented with aforementioned HMM profiles using hmmscan tool from HMMER v3.1b2 package and hits with an e-value of less than 10^−6^ were considered a match. Homologs of phi14:2 gp65 were found with the help of the jackhammer tool from the HMMER v3.1b2 package. Matching sequences had an e-value of less than 10^−6^. Two phage genomes (IAS^2^ and phicrAss001^27^) have been annotated previously. The putative promoter motifs were found more frequently upstream of phage genes encoding homologs of the phi14:2 middle proteins gp069 (function unknown), gp66 (RNAP), and gp074 (integration host factor IHF subunit), and late proteins gp092 (a structural protein of unknown function) and gp093 (portal) (**Supplementary file 1**). In 12 out of 14 phages the motif was at least once located upstream of a gene coding for tRNA. The DNA Logos of the motifs were constructed using WebLogo^37^.

### Purification of phi14:2 gp66 and *C. baltica* RNAP

The gene coding for the predicted phi14:2 RNAP catalytic subunit (g066 in this work; GeneID 16797463 in NCBI Reference Sequence NC_021806.1) was PCR amplified from phi14:2 genomic DNA and cloned into pETDuet-1 between BamHI and SacI restriction sites. This plasmid was used as a template to create mutant versions of g066 by site directed mutagenesis (list of corresponding primers is in **Supplementary Table 6**). Resulting plasmids were transformed into BL21 Star (DE3) chemically competent *E. coli* cells. The culture (3 L) was grown at 37°C to A_600_∼0.7 in LB medium supplemented with ampicillin at a concentration of 100 ug/mL and recombinant protein over-expression was induced with 1 mM IPTG for 3 hours at 20°C. Cells containing over-expressed recombinant protein were harvested by centrifugation and disrupted by sonication in buffer A (40mM Tris-HCl pH 8, 300mM NaCl, 3mM β-mercaptoetanol) followed by centrifugation at 15,000 g for 30 min. Cleared lysate was loaded onto a 5 mL HisTrap sepharose HP column (GE Healthcare) equilibrated with buffer A. The column was washed with buffer A supplemented with 20 mM Imidazole. The protein was eluted with a linear 0-0.5 M Imidazole gradient in buffer A. Fractions containing gp66 were combined and diluted with buffer B (40mM Tris HCl pH 8, 5% Glycerol, 0.5 mM EDTA, 1mM DTT) to the 50 mM NaCl final concentration and loaded on equilibrated 5 mL HiTrap Heparin HP sepharose column (GE Healthcare). The protein was eluted with a linear 0-1 M NaCl gradient in buffer B. Fractions containing gp66 were pooled and concentrated (Amicon Ultra-4 Centrifugal Filter Unit with Ultracel-30 membrane, EMD Millipore) to a final concentration 4 mg/ml, then glycerol was added up to 50% to the sample for storage at −20°C (the sample was used for transcription assays). For crystallization, fractions were diluted with buffer C (20 mM Tris HCl pH 8, 0.5 mM EDTA, 1mM DTT) to the 100 mM NaCl and loaded onto MonoQ 10/100 GL column (GE Healthcare). Bound proteins were eluted with a linear 0.1– 1 M NaCl gradient in buffer C. The fractions containing gp66 were pooled, diluted with buffer C to the 100 mM NaCl final concentration and concentrated to a final concentration 15 mg/mL and used for crystallization immediately.

To produce a Se-methionine (SeMet) derivative of gp66, the cells were first grown in the 2xTY medium until OD_600_ of 0.35, then pelleted by centrifugation at 4000 g for 10 min at 4°C and transferred to the SelenoMet Medium (Molecular Dimensions, Newmarket, Suffolk, UK) prepared according to the manufacturer’s instructions and supplemented with ampicillin at a concentration of 100 ug/mL. All the subsequent steps including the expression at low temperature and protein purification were the same as for the native protein.

For purification of *C. baltica* RNAP, 3 g of pelleted *C. baltica* cells were disrupted by sonication in 15 mL of buffer B (40 mM Tris-HCl pH 8.0, 0.5 mM EDTA, 1 mM DTT, 5% glycerol), containing 50 mM NaCl followed by centrifugation at 15,000 g for 30 min. Polyethylenimine P (pH 8.0) solution was added with stirring to the cleared lysate to the final concentration of 0.8 %. The resulting suspension was incubated on ice for 30 min and centrifuged at 10,000 g for 15 min. The pellet was washed by resuspension in buffer B with 0.3 M NaCl following centrifugation as previously. For elution, the pellet was resuspended in buffer B with 0.6 M NaCl. Eluted proteins were precipitated by adding ammonium sulfate to 67% saturation and centrifuged. The pellet was dissolved in 10 mL of buffer B and loaded onto a 1 mL HiTrap Heparin HP sepharose column (GE Healthcare) equilibrated with buffer B supplemented with 0.1 M NaCl. The column was washed with buffer B with 0.3 M NaCl, and RNAP was eluted with buffer B with 0.6 M NaCl. The fraction was concentrated by ultrafiltration (Amicon Ultra-4 Centrifugal Filter Unit with Ultracel-30 membrane, EMD Millipore) and loaded onto a Superdex 200 Increase 10/300 gel filtration column (GE Healthcare) equilibrated with buffer B containing 0.2 M NaCl. The fractions containing RNAP were pooled and concentrated up to 1 mg/mL, then glycerol was added up to 50% to the sample for storage at −20°C.

### DNA templates for transcription assay

For phi14:2 RNAP transcription assay genomic DNA of phi14:2 was purified using the Phage DNA Isolation Kit (Norgen Biotek Corp) according to the manufacturer’s instructions. Commercial genomic DNA of M13 bacteriophage (double- and single-stranded forms, New England Biolabs) were used.

For transcription by *C. baltica* RNAP, the PCR fragment containing T7 A1 promoter was used (5’ to 3’ sequence: tccagatcccgaaaatttatcaaaaagagtattgacttaaagtctaacctataggatacttacagcСatcgagagggccacggcgaa cagccaacccaatcgaacaggcctgctggtaatcgcaggcctttttatttggatccccgggta).

### *In vitro* transcription

Transcription reactions were performed in 5 µl of transcription buffer (20 mM Tris–HCl pH=8, 40 mM KCl, 10 mM MgCl_2_, 0.5 mM DTT and 100 µg/mL bovine serum albumin, RNAse inhibitor) and contained 100 nM gp66 and 50 ng genomic DNA. Where indicated, genomic DNA were denatured by heating to 100°C for 5 minutes following rapid cooling at 0°C. The reactions were incubated for 10 min at 22°C (or 10°C and 30°C where indicated), followed by the addition of 100 µM each of ATP, CTP, and GTP; 10 µM UTP and 3 µCi [α-^32^P]UTP (3000 Ci/mmol). Reactions proceeded for 30 min at 22°C (or 10°C and 30°C where indicated) and were terminated by the addition of an equal volume of denaturing loading buffer (95% formamide, 18 mM EDTA, 0.25% SDS, 0.025% xylene cyanol, 0.025% bromophenol blue). Where indicated, rifampicin was added to the final concentration of 50 µg/mL. Treatment with RNase T1 (Thermo Fisher Scientific) and DNAse RQ1 (Promega) were performed as follows: after the 30 min incubation of transcription reactions at 22°C, corresponding enzyme was added to 5 µl reactions; reactions were incubated for additional 15 min at 37°C and were terminated by the addition of an equal volume of denaturing loading buffer.

The reaction products were resolved by electrophoresis on 5 % (w/v) denaturing 8 M urea polyacrylamide gel. Since high-molecular weight RNA was expected to be synthesized from genomic DNA templates, the electrophoresis was run for 2 hours. Transcription reaction products by *C. baltica* RNAP were loaded on the gel with a delay to observe both, high-molecular weight RNA synthetized by gp66 from genomic DNA and 67 nucleotides RNA synthesized by *C. baltica* RNAP from PCR fragment. Results were visualized by Typhoon FLA scanner (GE Healthcare).

Transcription reactions from RNA-DNA scaffolds were set at the same buffer as above transcription reactions and contained 15 nM RNA-DNA scaffold and 15 nM of gp66, T7 RNAP or *E. coli* RNAP core (New England Biolabs). Reactions were incubated for 10 min at 22°C, followed by the addition of 1mM each of ATP, CTP, GTP and UTP. Reactions proceeded for 30 min at 22°C and were terminated by the addition of an equal volume of denaturing loading buffer; the products were resolved by electrophoresis on 16 % (w/v) denaturing 8 M urea polyacrylamide gel. Results were visualized by Typhoon FLA scanner (GE Healthcare).

All transcription experiments were repeated at least three times.

### Crystallization and structure determination of phi14:2 RNAP (gp66)

The initial crystallization screening was carried out by the sitting drop method in 96 well ARI Intelliwell-2 LR plates using Jena Bioscience crystallization screens at 19°C. PHOENIX pipetting robot (Art Robbins Instruments, USA) was employed for preparing crystallization plates and setting up drops each containing 200 nl of the protein and the same volume of the well solution. Optimization of crystallization conditions was performed in 24 well VDX plates and thin siliconized cover slides (both from Hampton Research) by hanging drop vapor diffusion. Crystallization drops of the 24-well plate setup contained 1.5 μl of the protein solution in 20 mM Tris-HCl pH 8.0, 100 mM NaCl, 1 mM DTT, 0.5 mM EDTA mixed with an equal volume of the well solution. Best crystals of both native and SeMet gp66 were obtained with the protein having the initial concentration of 15 mg/ml and equilibrated against 700 μl of the well solution containing 100 mM Tris-HCl pH 8.5, 200 mM NaOAc, 11% PEG 4000, 2 mM TCEP. Ta_6_Br_12_ derivatized crystals of gp66 were produced by soaking native crystals in a pre-equilibrated crystallization solution that contained a freshly prepared Ta_6_Br_12_ compound at a 1-2 mM concentration. Upon soaking for 1-3 days, Ta_6_Br_12_ derivatized crystals acquired an emerald green color. For data collection, the crystals were dipped for 15 seconds into cryo solutions containing 30% of glycerol in addition to the well solution components and flash frozen in liquid nitrogen. X-ray diffraction data and fluorescent spectra were collected in a nitrogen stream at 100 K.

X-ray fluorescence emission spectra of both Ta_6_Br_12_ and SeMet derivative crystals displayed a strong “white line” at the L_III_ and K absorption edges of Ta and Se, respectively. The corresponding excitation wavelengths – 1.25478 Å for Ta_6_Br_12_ crystals and 0.97872 Å for SeMet – were then used for data collection. Diffraction data were collected on two different beamlines of the Life Sciences Collaborative Access Team at Advanced Photon Source, Chicago: Ta_6_Br_12_ on 21-ID-D (Dectris Eiger 9M area detector), and SeMet on 21-ID-F (Rayonix MX300 area detector). All datasets comprised a full 360° swath that was cut into 0.125° frames on the Eiger detector (2880 frames) or 0.5° frames on the Rayonix detector (720 frames). The datasets were indexed, integrated, and reduced with the help of the XDS suite. The heavy atom substructure of Ta_6_Br_12_ datasets that had an anomalous signal greater than 1.37 (as defined by XDS) could be easily solved. On the other hand, all attempts at *ab initio* solution of the Se atom substructure in SeMet datasets with an anomalous signal as great as 1.28 failed. The Se substructure was expected to consist of around 80 atoms because the asymmetric unit contained two molecules of gp66 with 39 methionines each.

An interpretable electron density was obtained as follows. First, we solved the heavy atom substructure of one of the Ta_6_Br_12_ soaked dataset with the help of HKL2MAP and SHELXD suite. These phases were improved by two-fold non-crystallographic averaging. A large fraction of the polypeptide chain could be traced in this electron density, but it had a resolution of 3.75 Å and it was discontinuous and disordered in places. The partial model was then used in a molecular replacement procedure to solve the best SeMet dataset. The height of the peaks in the Bijvoet difference Fourier synthesis map of the SeMet dataset decreased gradually, and the exact number of ordered Se sites could not be established. For this reason, the first 71 peaks that were somewhat higher than the rest were input into PHASER to find all Se sites and obtain new phases. These phases were improved by two-fold non-crystallographic averaging with the help of the program PARROT. The resulting 3.5 Å resolution electron density could be traced with relative ease and interpreted in nearly all 2,180 residues comprising gp66 barring for a few residues at both termini.

The atomic model was refined with the help of the programs PHENIX and COOT. Molprobity was used in the validation procedure. The final model has 94.76% of residues in the favorable region of the Ramachandran plot and 0.07% outliers.

## Data availability

Genome of *C. baltica* strain 14 has been deposited in the NCBI BioProject and is accessible through BioProject ID PRJNA552277.

The RNA-Sequencing data have been deposited in the NCBI Gene Expression Omnibus^38^ and are accessible through GEO Series GenBank accession no. GSE133609.

The refined atomic model of phi14:2 gp66 and the associated experimental data have been deposited to the Protein Data Bank under the accession number 6VR4.

Additional data are available from the corresponding authors upon request.

## Acknowledgments

We would like to thank Sofia Medvedeva (Skolkovo Institute of Science and Technology, Moscow, Russia) for help with promoter search. The study was carried out using resources of the Skoltech Genomics Core Facility. The work was supported by the Russian Science Foundation (grant no 19-74-00011 to M. L. Sokolova).

## Author contributions

**K.V.S**., **M.L.S**. and **E.V.K** conceived the study. **K.H**. and **E.N**. provided *C. baltica* cells, phi14:2 phage and phi14:2 DNA. **A.V.D**. cultivated *C. baltica* and phi14:2, prepared RNA for RNA-Seq and primer extension experiments (PE), performed RT-qPCR. **S.P**. purified phi14:2 RNAP and its mutants, performed all *in vitro* transcription assays and some of the PE. **M.K**. processed and analyzed RNA-Seq data, annotated crAss-like phage genomes. **E.I.K**. performed mutagenesis of phi14:2 RNAP. **L.M**. performed PE. **M.V.Y**. purified *C. baltica* RNAP. **M.L.S**. performed search for promoters, prepared crystals. **P.G.L**. solved crystal structure. **M.L.S**., **P.G.L**., **S.B**. analyzed the structure. **M.L.S**., **P.G.L**., **K.V.S**. wrote the manuscript, which was read, edited and approved by all authors.

---

**Extended data Fig. 1.**
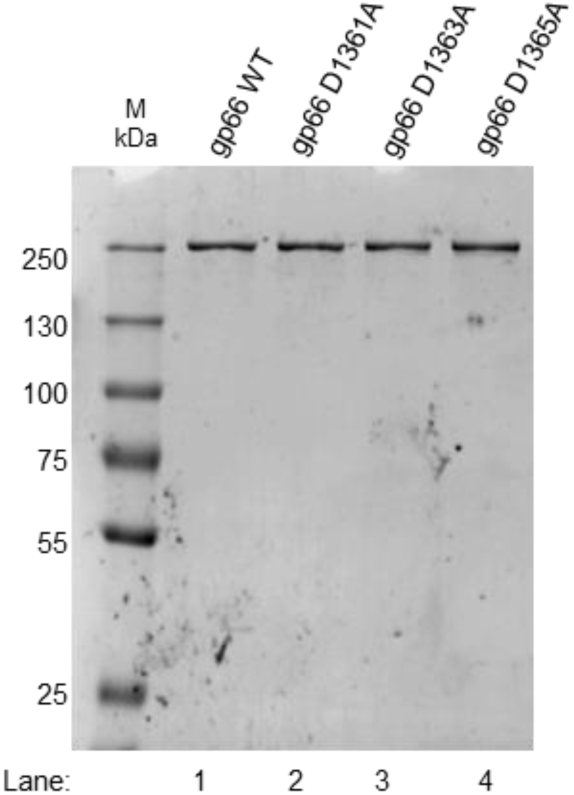
SDS-PAGE analysis of wild-type gp66 and mutant gp66. SDS-PAGE analysis of wild-type gp66 and gp66 mutants carrying single alanine substitutions of each aspartate in the DFDID motif purified by Heparin HP sepharose column chromatography.

**Extended Data Fig. 2.**
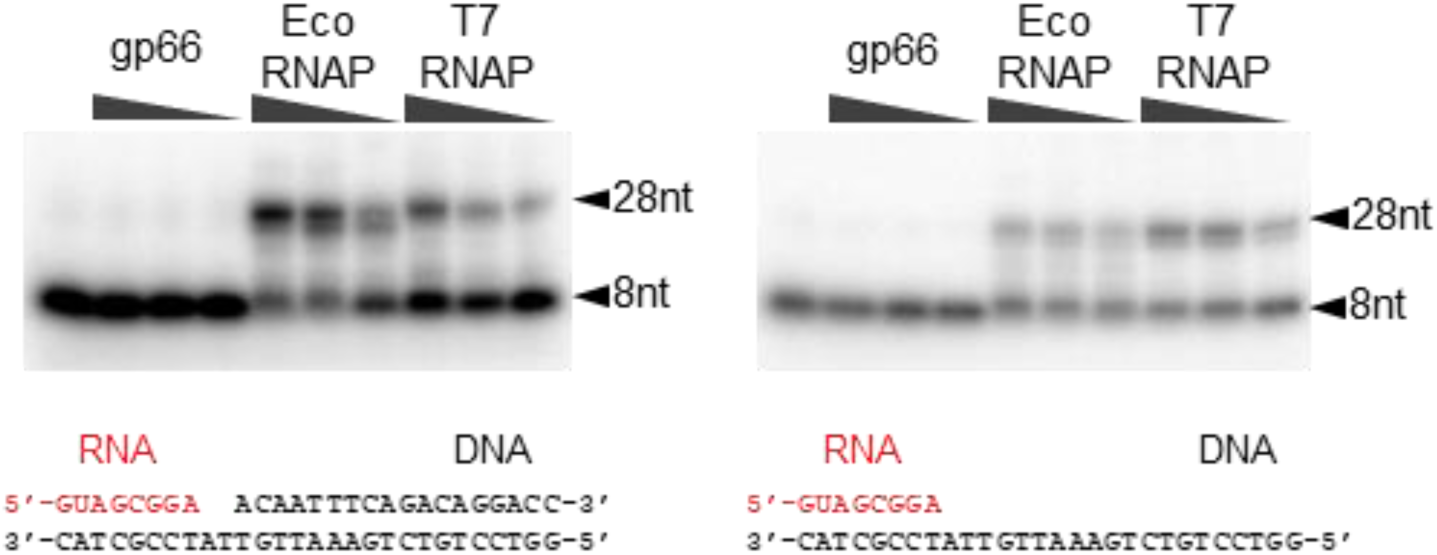
Gp66 does not extend RNA primer in RNA-DNA scaffold. Extension of RNA primer in RNA-DNA scaffolds by gp66, *E. coli* (Eco) and T7 RNAPs as controls in the presence of ribonucleoside tri-phosphates. The sequences of RNA-DNA scaffolds used are shown under the gels; the RNA was radioactively labeled at the 5′ end. The reaction products were resolved by electrophoresis in 16 % (w/v) denaturing 8 M urea polyacrylamide gel and revealed by autoradiography.

**Extended Data Fig. 3.**
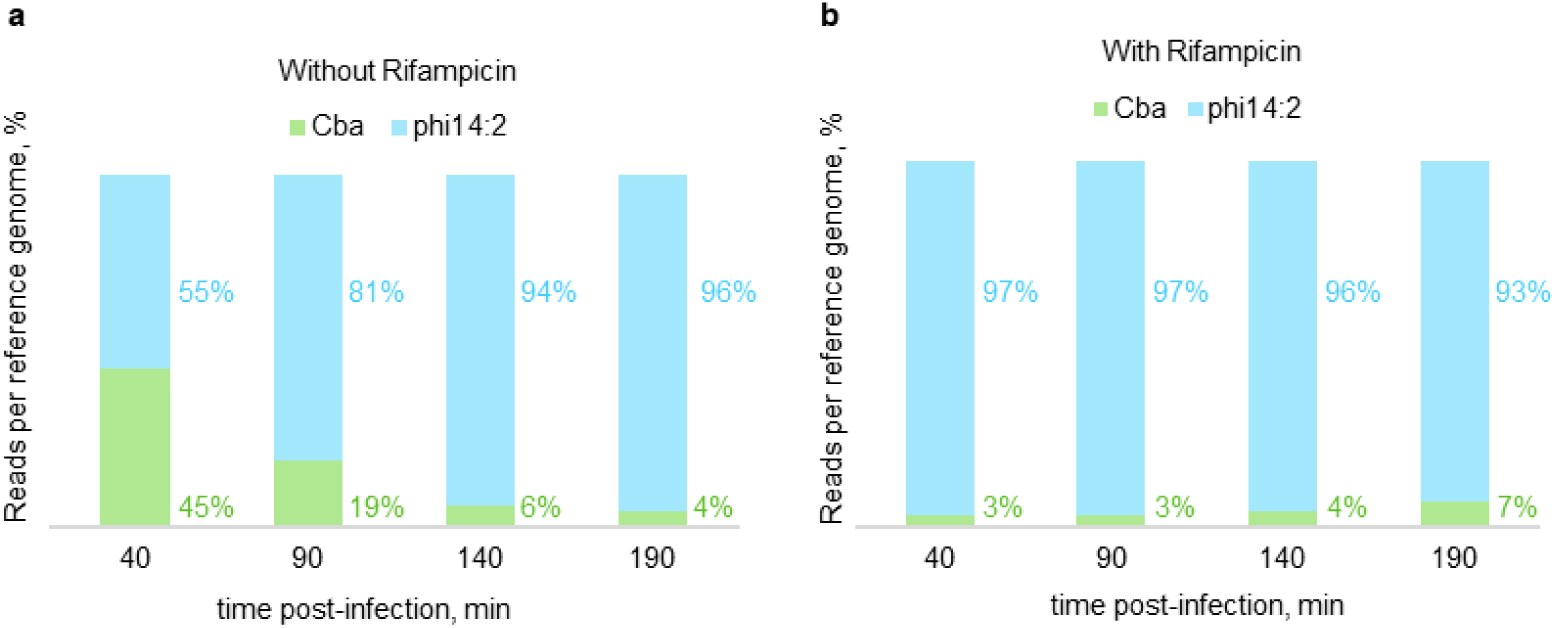
Distribution of phage and host transcript abundances. The total number of reads per reference sequence aligned with a corresponding genome (Cba – *C. baltica* strain 14, this study; phi14:2 – NC_021806) is shown for the Rif-libraries (**a**) and for the Rif+ libraries as stacked bars (**b**). The percentages are indicated next to the bars.

**Extended Data Fig. 4.**
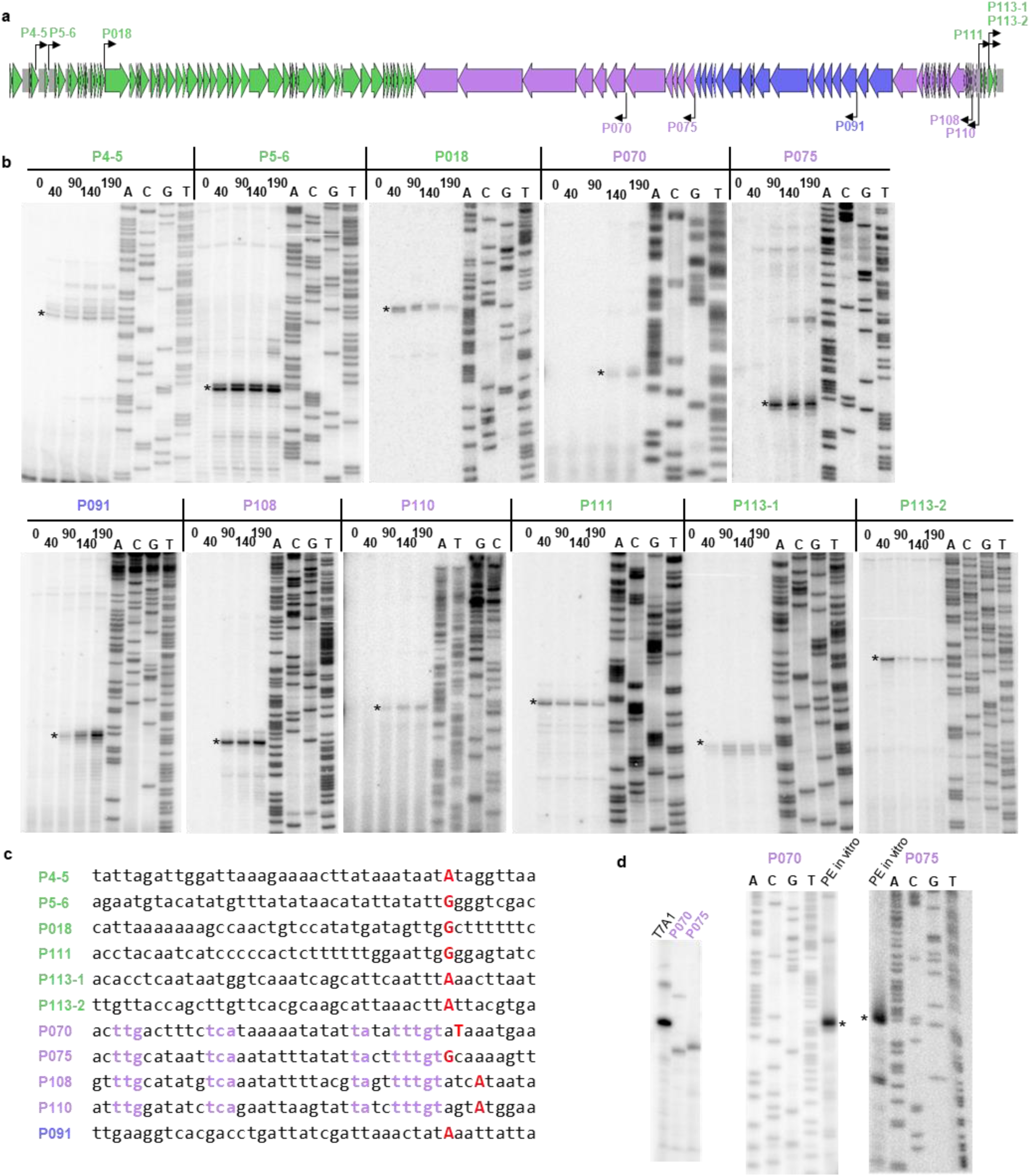
Identification of putative promoters in phi14:2 genome. **a**, Schematic of the phi14:2 genome (see the legend of Fig. 2a for details) with putative promoters marked by black arrows. **b**, Primer extension and sequencing reactions for eleven putative promoters (Supplementary Table 4). Major primer extension products are marked with black asterisks. **c**, Sequences flanking the primer extension endpoints are shown; nucleotides, corresponding to primer extension endpoints are colored red. Conserved nucleotides of putative middle promoters are shown in violet. **d**, Left panel: *In vitro* transcription of PCR-fragments containing the T7 A1 promoter and predicted phi14:2 P070 and P075 promoters by *E. coli* RNAP; Right panel: Primer extension reactions of RNA synthesized *in vitro* by *E. coli* RNAP from PCR-fragments containing phi14:2 P070 and P075 promoters.

**Extended Data Fig. 5.**
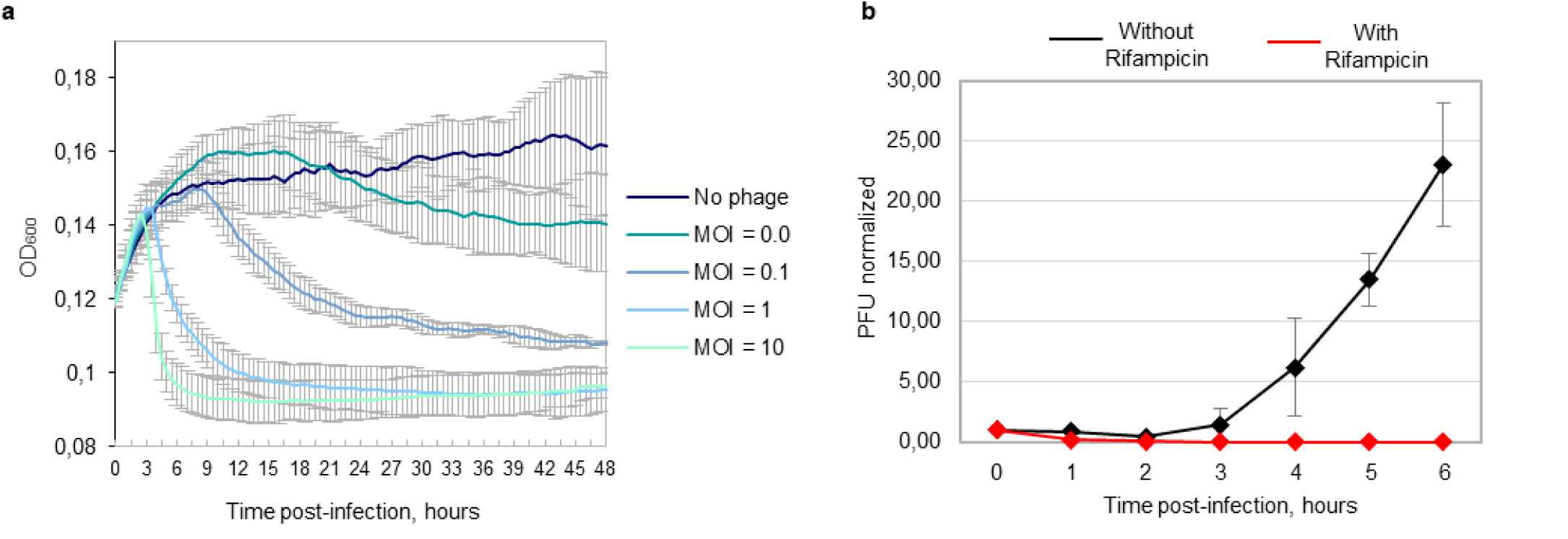
General parameters of phi14:2 infection. **a**, Growth curves of *C. baltica* infected with phi14:2 at different MOIs in the log growth phase (mean±SD of three biological replicates). **b**, Single-burst curves of phi14:2 infecting *C. baltica* at MOI∼0.5. Number of plaque forming units (PFUs) normalized to the PFU immediately after the phage was added to the culture (0 time point) (mean±SD of three biological replicas) are shown for cultures treated (red line) or not treated (black line) with host RNAP inhibitor rifampicin (Rif) prior to infection.

**Extended Data Fig. 6.**
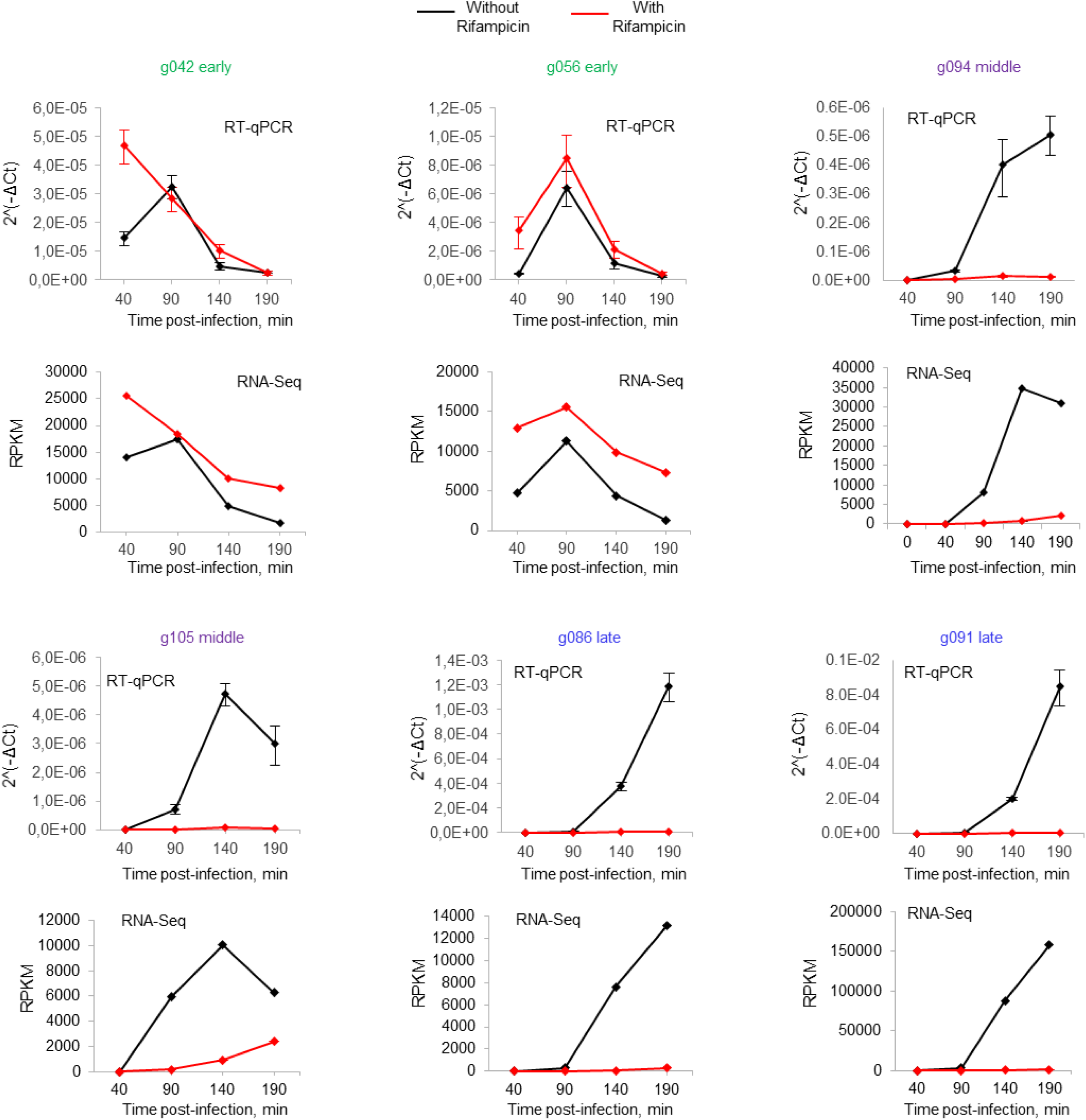
Validation of RNA-Seq data by RT-qPCR. Relative transcript abundances of six selected phi14:2 genes during the infection of *C. baltica* cells in the presence (red) and absence (black) of rifampicin were determined by RT-qPCR. The cycle threshold (Ct) values of the *C. baltica* 16S RNA were used to normalize the Ct values of selected phi14:2 transcripts as follows: ΔCt = (mean Ct gene) – (mean Ct 16S rRNA)). The amplicon concentrations for different time points are plotted as 2^(−ΔCt) (mean±SD of three technical replicates). Corresponding RNA-Seq data are shown below the results of RT-qPCR.

**Extended data Table 1.** Absolutely conserved amino acids of RNAPs of crAss-like phages and their analogs in other RNAPs based on structural alignments. Residue numbers are given for gp66 of phi14:2, QDE-1 RNAP of *Neurospora crassa* and RNAP of *Thermus thermophilus* (T. th). Light green-colored cells describe amino acids conserved in all three types of RNAPs; dark green-colored cells indicate amino acids conserved in crAss-like phage and multisubunit RNAPs; dark blue-colored cells show amino acids conserved in crAss-like phage RNAPs and QDE-1 RNAP; light blue-colored cells contain amino acids unique to crAss-like phage RNAPs.

| No | *phi14:2*<br>*gp66* | *QDE-1* | *T. th* | **Function in canonical DNA-dependent RNAPs according to analysis by Lane and Darst<sup>20</sup>**<br>Residue numbers are indicated for *T. th* RNAP |
| --- | --- | --- | --- | --- |
| 1 | Lys893 | - | - | - |
| 2 | Arg894 | Arg671 | β-Arg557 | β-Arg557 interacts with the γ-phosphate of the rNTP; |
| 3 | Asp962 | Asp709 | β-Asp686 | β-Asp686 interacts with: 1) β'-Asp739/Phe740/Asp741 of the β'-NADFDGD motif; 2) the γ-phosphate of rNTP and MgII in the active site; 3) β-Arg879, which also interacts with the rNTP γ-phosphate; |
| 4 | Lys1012 | - | β-Lys838 | β-Lys838 and β-Lys846 interact with the backbone of the RNA transcript at the -1/-2 positions; |
| 5 | Lys1027 | Lys743 | β-Lys846 |  |
| 6 | Lys1065 | Lys767 | - | - |
| 7 | Gln1116 | Gln797 | - | - |
| 8 | Gly1235 | - | - | - |
| 9 | Arg1322 | Arg962 | β'-Arg704 | β'-Arg704 interacts simultaneously with the O4' of rNTP, the 2'-OH of the RNA transcript at -1 position, and β'-Asn737, β'-Ala738, and β'-Asp743 of the β'-NADFDGD motif; |
| 10 | Pro1324 | Pro964 | β'-Pro706 | β'-Pro706 lines the path for the template DNA around the +1 position; |
| 11 | Ser1330 | - | - | - |
| 12 | Gly1359 | Gly1005 | β'-Asn737 | Within the β'-NADFDGD motif, β'-Asn737 interacts with O2' and O3' of the rNTP; |
| 13 | Asp1361 | Asp1007 | β'-Asp739 | β'-Asp739 interacts with Mgl and MgII, and with absolutely conserved β-Asp686; |
| 14 | Asp1363 | Asp1009 | β'-Asp741 | β'-Asp741 interacts with Mgl, the RNA transcript at the -1 position, and β-Asp686; |
| 15 | Asp1365 | Asp1011 | β'-Asp743 | β'-Asp743 interacts with Mgl and the RNA transcript at the -1 position. |
| 16 | Asp1612 | Asp1116 | - | - |
| 17 | Lys1615 | Lys1119 | - | - |

